# Pain and the Emotional Brain: Affective Rather than Cognitive Processes Drive the Cortical Encoding of Pain

**DOI:** 10.1101/2021.06.30.450586

**Authors:** Anne Stankewitz, Astrid Mayr, Stephanie Irving, Viktor Witkovsky, Enrico Schulz

**Affiliations:** Department of Neuroradiology, Technische Universität München, Munich, Germany; Department of Radiology, LMU University Hospital, Ludwig-Maximilians-Universität München, Munich, Germany; Department of Neurology, LMU University Hospital, Ludwig-Maximilians-Universität München, Munich, Germany; Department of Theoretical Methods, Institute of Measurement Science, Slovak Academy of Sciences, Bratislava, Slovak Republic; Wellcome Centre for Integrative Neuroimaging, FMRIB, Nuffield Department of Clinical Neurosciences, University of Oxford, Oxford, UK; Department of Medical Psychology, Ludwig-Maximilians-Universität München, Munich, Germany

**Keywords:** pain, fmri, unpleasantness, 7 tesla

## Abstract

**Background:** The experience of pain has been dissociated into two interwoven aspects: a sensory-discriminative aspect and an affective-motivational aspect. We aimed to explore which of the pain descriptors is more deeply rooted in the human brain.

**Findings:** Participants were asked to evaluate applied cold pain. The majority of the trials showed distinct ratings: some were rated higher for unpleasantness and others for intensity. We compared the relationship between functional data recorded from 7 tesla MRI with unpleasantness and intensity ratings and revealed a stronger relationship between cortical data and unpleasantness ratings.

**Conclusions:** The present study underlines the importance of the emotional-affective aspects of pain-related cortical processes in the brain. The findings corroborate previous studies showing a higher sensitivity to pain unpleasantness compared to ratings of pain intensity. For the processing of pain in healthy subjects, this effect may reflect the more direct and intuitive evaluation of emotional aspects of the pain system, which is to prevent harm and to preserve the physical integrity of the body.

## 1. INTRODUCTION

The experience of pain has been dissociated into sensory-discriminative and affective-motivational aspects [1, 2], which are widely assessed as subjective ratings of pain intensity and pain unpleasantness [3–5]. Both descriptors of pain share a conceptual core variance and are therefore often highly correlated: very intense pain is associated with higher levels of unpleasantness [6, 7]. By contrast, previous research also emphasised the disparity of both pain descriptors by revealing a diverging evaluation depending on the type of pain, for example health-related pain conditions are associated with higher pain unpleasantness ratings [2]. Differential ratings of intensity and unpleasantness have been revealed in dependence on the type of experimentally-applied pain [8]. Likewise, the experience of pain unpleasantness is considered to exhibit a particularly strong influence on well-being [9, 10], as it has been found to be tightly connected to pain catastrophising [9].

Studies exploring the cortical underpinnings of pain unpleasantness are scarce and non-specific as they implicitly include the shared variance with pain intensity [11, 12]. The conceptual and empirical proximity of both aspects has made an investigation on the differences between the cortical underpinnings of pain intensity and pain unpleasantness challenging. The only attempt to experimentally dissociate pain intensity and pain unpleasantness processing has been conducted in a series of early PET studies. Using hypnosis, a specific modulation of pain unpleasantness was reported in the anterior cingulate cortex (ACC) but not for the somatosensory cortices (SI and SII) and the insular cortex (IC) [13]. This clear effect was shadowed by a subsequent study [7], where the authors were not able to hypnotically disentangle intensity and unpleasantness. The ACC activation was slightly shifted compared to the previous study and despite (unintended) differences in unpleasantness in response to the hypnotic modulation of pain, there was no effect for the ACC. The shift of activity in the ACC region between studies, the absence of information on the sample size for one study [13], and the absence of an ACC effect in the follow-up study leave some questions regarding distinct cortical processing of pain intensity and pain affect in this region.

Nonetheless, despite a high correlation between both subjective pain aspects, it is indeed possible to dissociate the cortical underpinnings of the encoding of pain intensity and pain unpleasantness in a within-subject design without the need for any potentially unreliable modulatory intervention. To this end, we aimed to investigate the brain regions that differentially process pain intensity and pain unpleasantness. As a major advantage, by avoiding undesirable session or sample variability, both aspects are explored in relation to the very same underlying cortical data; each trial has been evaluated for either descriptor of the subjective experience of pain. The majority of the trials indeed differed and enabled us to elucidate which aspect of pain perception is more deeply rooted in the human brain.

## 2. MATERIALS AND METHODS

### Subjects

20 healthy subjects (16 female/4 male, age 27 ± 5 years; mean ± standard deviation) were included in the study. All subjects gave written informed consent, none reported any history of chronic pain.

The data were taken from our previous study and the experimental procedure has been described in detail in our previous publications [14, 15]. The previous studies explored the cortical underpinnings of cognitive interventions to attenuate pain: the experiment consisted of four conditions across four separate blocks, where each block comprised 12 trials from the same condition. In all conditions and trials, the subjects received cold pain stimuli on the dorsum of their left hand delivered by a thermode (Pathway II; Medoc Ltd, Israel). The first condition of the experiment was always the unmodulated pain condition. The subsequent three conditions were counterbalanced blocks of pain attenuation: (A) an attentional shift, (B) an imaginal strategy, and (C) a non-imaginal reinterpretation. After each trial, the subjects were prompted to rate pain intensity and pain unpleasantness within 10 s (Figure 1). The scale ranged between 0 and 100 in steps of 5 points. The endpoints of the scale were determined as no pain (0) and the maximum pain the subjects were willing to tolerate (100). There were a total of 48 trials during the fMRI recording. The 40 s of painful stimulation were preceded by a rest period of 10 s at 38 °C thermode temperature. The different conditions and their modulation of pain across trials are not relevant here as we are exploring the within-trial differences between two descriptors of pain in the same trial. Our approach avoids a distinct modulating intervention for intensity or unpleasantness, which may not always work as intended [7]. In addition, a distinct intervention for either descriptor of pain may introduce undesirable effects of variability due to recordings from different sessions or samples.

**Figure 1.**
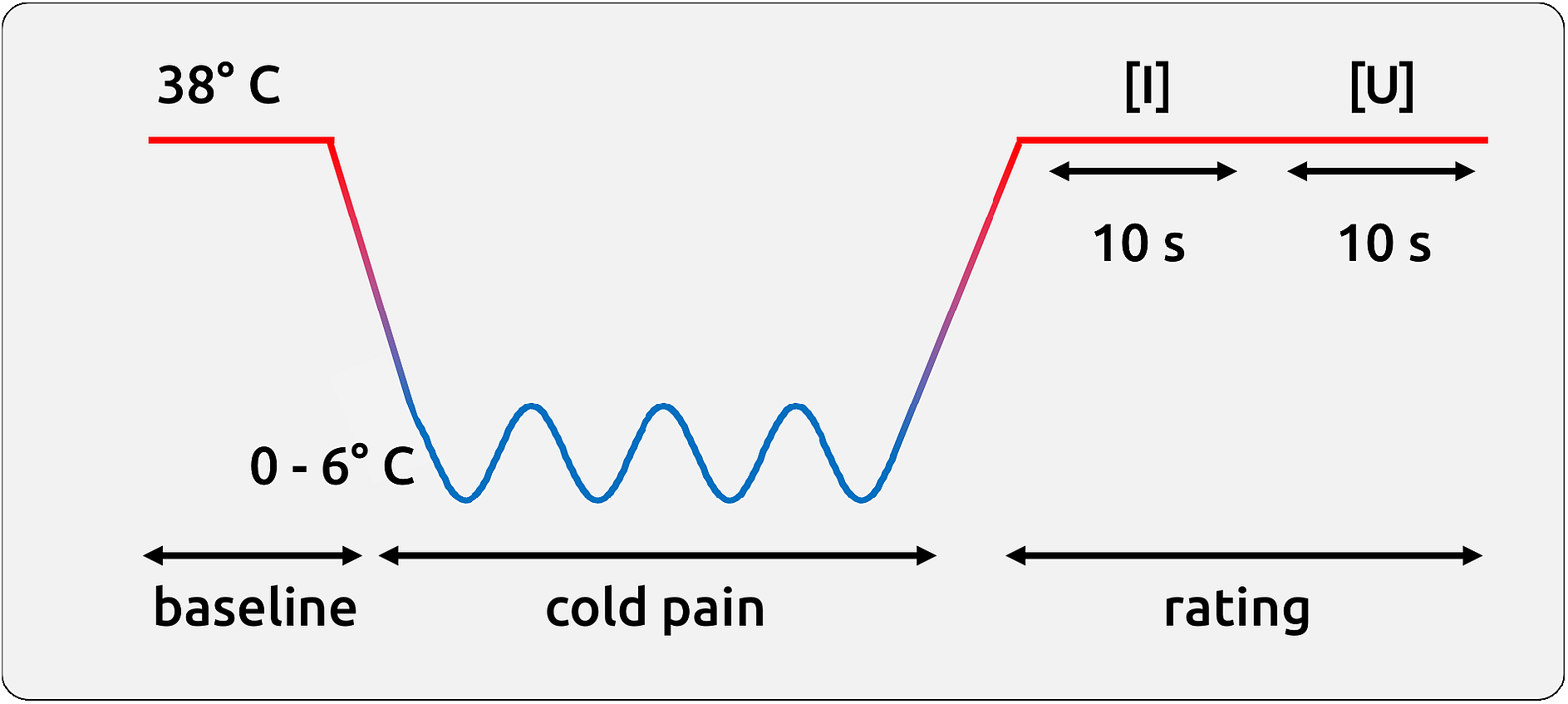
Schematic time course of a single trial. After 10 s of rest, painful stimulation was applied for 40 s. The pain was rated regarding intensity ([I], 10 s) and unpleasantness ([U], 10 s).

### Data Acquisition

Imaging data were acquired on a 7 tesla Siemens MRI scanner using parallel image acquisition (GRAPPA, factor = 2). In order to cover the whole brain, each of the 1768 functional echo-planar imaging (EPI) volumes comprised 34 axial slices of 2 mm thickness and 2 × 2 mm in-plane resolution with a 1 mm gap between slices. The repetition time (TR) was 1.96 s, the echo time (TE) was 25 ms (flip angle 90°), the field of view (FOV) was 220 × 220 mm, and the matrix size was 110 × 110 pixels. A T1-weighted structural image (isotropic 1 mm^3^ voxel) was acquired for the registration of the EPI to the MNI (Montreal Neurological Institute) template.

### Imaging analysis - preprocessing

The data were pre-processed and analysed with FSL [16] and Matlab (version R2020a, The Mathworks, USA). The preprocessing consisted of brain extraction, high-pass filtering with a frequency cutoff of 1/90 Hz, a correction for head motion during scanning, spatial normalisation to the MNI template, and spatial smoothing (6 mm FWHM). The data were further semi-automatically cleaned of artefacts with MELODIC [17]. Beta coefficients representing the magnitude of cortical activity for each trial (modulated and unmodulated) were computed in FEAT [15].

### Image analysis - extraction of regions of interest data

The time series of functional volumes were converted to MNI space and subsequently projected to surface space by using the “Connectome Workbench” package. We used a template that allowed us to project from 3D standard MNI space to 2D surface space. Regions of interest (ROIs) were defined by subdividing the cortical surface into 180 regions per hemisphere [18]. Six further regions (5 bilateral) that are important for the processing of pain, such as the PAG, the thalamus and the amygdala, were also included. The ROIs were based on the Oxford Atlas, implemented in FSL.

### Image analysis - computation of single trial functional connectivity scores

The time courses for all voxels of cortical activity for a specific region of the Glasser Atlas, e.g. the anterior insula, were extracted. We computed principal component analyses (PCA) separately for each ROI and subject and selected the first component (Matlab, The MathWorks, Inc., USA). The plateau phase of the last ~30s of painful stimulation (15 data points), derived from the FEAT design matrix, has been extracted from each region and trial for each subject and condition. Outliers were removed from the data by using the Grubbs’ test (Grubbs, 1950). These 15 data points determined the connectivity for a brain region for a given trial. Kendall’s tau correlation coefficients (*τ*) were computed for each trial and for each of the 371 ROIs with the remaining 370 ROIs. The single trial correlation coefficients were Fisher Z-transformed and fed into group-level statistical analysis.

### Statistical analysis - comparison between intensity and unpleasantness

Differences between pain intensity and pain unpleasantness occurred in 75% of the trials. Using Linear Mixed Effects models [19, 20], we aimed to differentiate the relationship between the two aspects of pain ratings and cortical activity. The statistical model is expressed in Wilkinson notation [21]; the included fixed effect of interest (rating_type:fmri) describes the magnitudes of the population common intercept and the population common slopes for the relationship between cortical data and the single-trial differences in pain perception. The added random effect (i.e. 1|subject) models the specific intercept differences for each subject (e.g. subject-specific differences in fMRI beta coefficients):

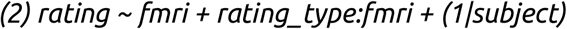

In this formula, *rating* concatenates the single trial pain ratings (unpleasantness, intensity) and *fmri* relates to the single trial estimate of the BOLD response, which is identical for the corresponding ratings of intensity and unpleasantness (see example in Supplementary Material). Consequently, for each voxel, the statistics include all single-trial pain intensity ratings and the respective cortical data, as well as all single-trial unpleasantness ratings and the same cortical data. Each *rating* entry was assigned its corresponding single-trial grouping code *(rating_type, nominal[-1 1])* and BOLD estimate (*fmrl*). Please note our more suitable modelling in Supplementary Material, where we - by generating a gradient vector - put more weight on trials that had greater single-trial rating differences.

Due to its relative character, we can not know whether a potential negative difference is due to a weaker positive relationship or a stronger negative relationship. Therefore, the results can only be evaluated in light of the general map of pain processing (activation, deactivation). Consequently, we created a 3D map consisting of the group result of the analysis in FEAT by contrasting the pain trials with the no-pain baseline period (cluster corrected, p<0.05). The LME analysis was restricted to the significant voxels within this map. As delineated above, the direction of the contrast (activations, deactivations) determines the interpretation of the direction of effect for the differential encoding of pain intensity and pain unpleasantness.

To correct for multiple comparisons, we applied a randomisation approach. Behavioural data were shuffled and the entire analysis was repeated 5000 times within the boundaries of the 3D brain mask. The highest absolute t-values of each repetition across the whole volume were extracted. This procedure resulted in right-skewed distribution for each condition. Based on these distributions, the statistical thresholds were determined using the “palm_datapval” function publicly available in PALM [22].

### Statistical analysis - mapping of pain processing

In order to interpret the relative findings between pain intensity and pain unpleasantness it was required to assess whether cortical processes increase or decrease in response to pain. For the BOLD data, we contrasted in FEAT the phases of pain experience (all 4 conditions pooled) compared to the baseline period. The map closely resembles the neurologic signature for pain map [23]. For connectivity data, we also utilised the information from the design matrix and compared the pooled connectivity of all pain phases with the connectivity in baseline periods. The majority of the pain trials (97%) and the baseline periods (98%) showed a positive ***τ*** at single trial level. Similar to the interpretation of the BOLD data, the results of these analyses are relevant insofar as we can determine whether a lower connectivity for unpleasantness can be interpreted as weaker coupling (in case we observe general increase of connectivity during pain compared to the baseline) or as stronger anticorrelation (in case we observe general decrease of connectivity during pain compared to the baseline). For BOLD and connectivity analyses, we will mask the comparison results (intensity vs. unpleasantness) with the findings from the pain mapping (pain vs. baseline) using a liberal threshold (abs(t)>2).

## 3. RESULTS

### 3.1. Behavioural data

The single-trial pain ratings indicate that the applied cold pain is particularly suited for distinguishing the cortical underpinnings of pain intensity and pain unpleasantness. Unlike previous research [8], we did not find any systematic rating difference between both descriptors (paired t-test, p<0.05; 34 ±14 for pain intensity and 31±17 for pain unpleasantness). Although highly correlated (r=0.86, p<0.01, Figure 2), the majority of the trials exhibited a distinct rating for intensity and unpleasantness (75%, Figure 2). Only 25% of the trials were rated identically (e.g. 25/100 for intensity and 25/100 for unpleasantness). The trial-by-trial differences in pain ratings can be utilised to elucidate whether the cortical activity is more closely related to pain intensity or to pain unpleasantness.

**Figure 2.**
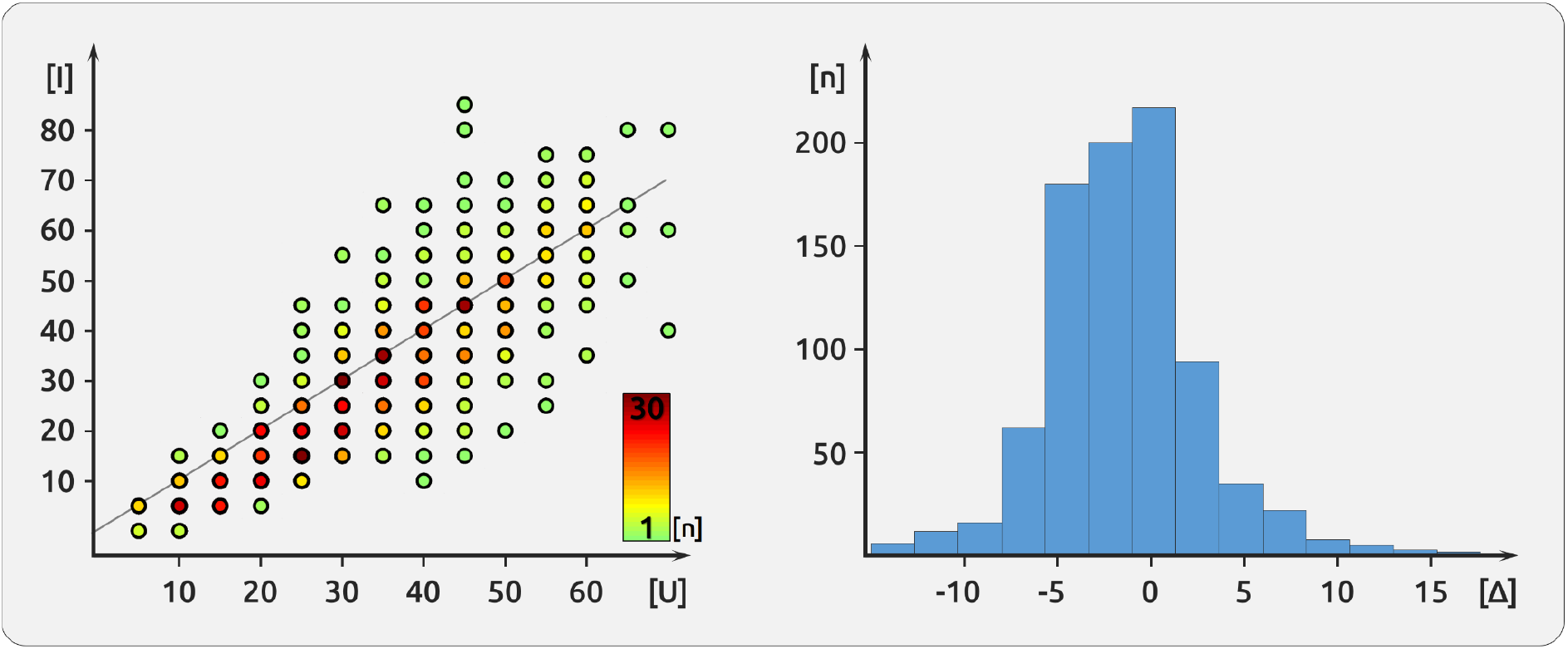
Description of pain ratings. The left side (A) shows the correlation between pain intensity [I] and pain unpleasantness [U]. The correlation coefficient is r=0.86. Nevertheless, the ratings for both aspects were identical only for a minority of the trials. The colour-coded points indicate the frequency of the occurrence of each single-trial combination of intensity and unpleasantness ratings. Near-identical ratings of intensity and unpleasantness occur more often than more distant ratings. The histogram (B) shows the difference between pain intensity ratings and pain unpleasantness ratings; most trials (75%) exhibited different ratings. For some trials, pain intensity was rated higher (negative Δ), for others pain unpleasantness was rated higher (positive Δ).

### 3.2. Imaging data

#### Positive effects - BOLD

The results indicate a stronger positive relationship for *pain unpleasantness* in regions that are known to exhibit higher activity in response to experimental pain: the bilateral insula and frontal opercular cortex, pre- and postcentral regions, as well as in frontal regions (p<0.05, PALM corrected). These positive effects occur exclusively in brain regions that are activated in response to pain in the contrast of the unmodulated pain trials compared to the baseline (pale colours in Figure 3).

**Figure 3.**
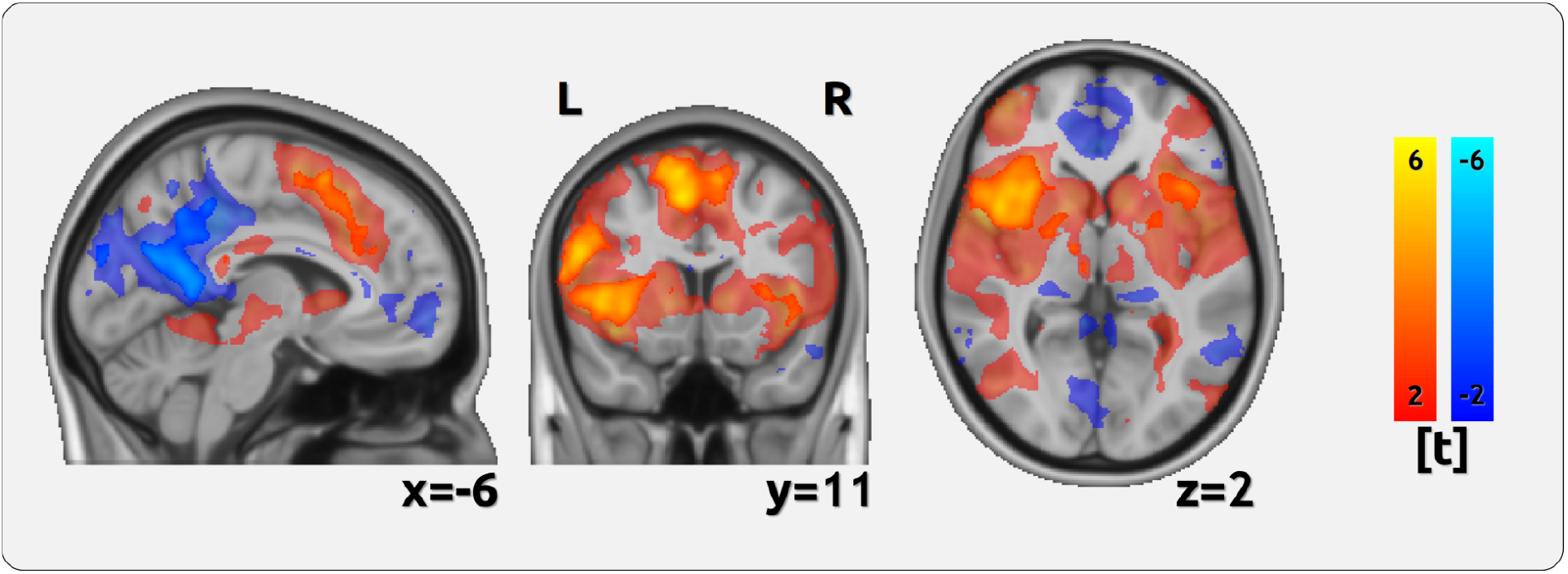
Differences between pain unpleasantness encoding and pain intensity encoding - BOLD. Several cortical regions reflect the differences between pain intensity and pain unpleasantness. Warm colours indicate regions that show a more positive relationship for pain unpleasantness; cold colours indicate brain regions that show a stronger negative relationship for pain unpleasantness. The pale colours indicate the contrast between pain trials and baseline period (abs(t)>2).

#### Negative Effects - BOLD

Additionally, we found a more positive relationship for *pain intensity* for cuneal and precuneal regions, the occipital cortex, as well as angular and frontal regions (p<0.05, PALM corrected). These negative results occur exclusively in brain regions that are suppressed in response to pain in the contrast of the unmodulated pain trials compared to the baseline (pale colours in Figure 3). Therefore, they reflect a stronger negative relationship between pain ratings and brain activity for unpleasantness (Figure 3, Table 1).

**Table 1.**
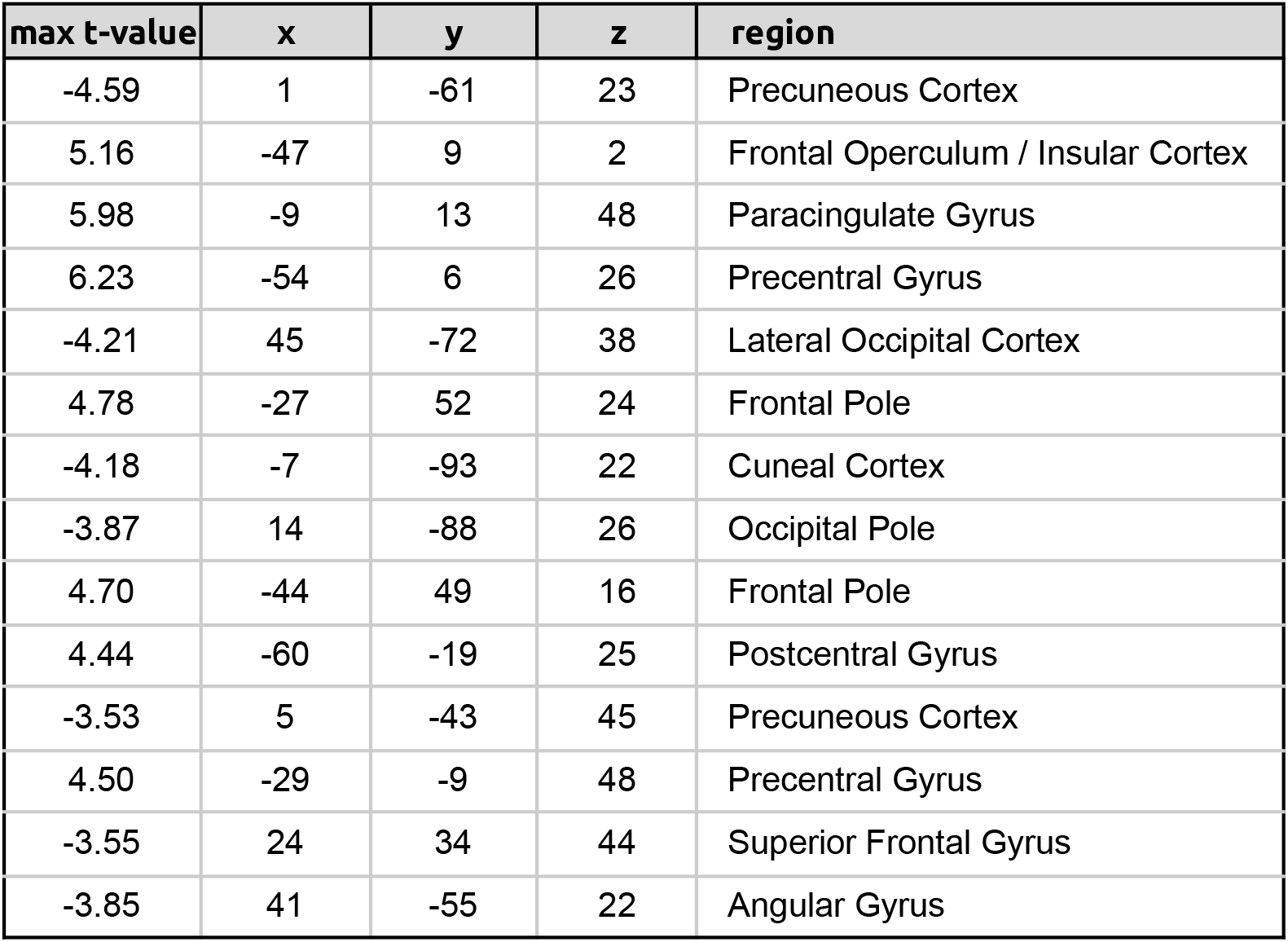
Differences in the encoding of pain intensity and pain unpleasantness. Positive effects indicate a stronger positive relationship for unpleasantness; negative relationships indicate a stronger negative relationship for unpleasantness (all p<0.05, PALM corrected).

#### Positive effects - connectivity

We did not find any positive effects for two reasons. The comparison for pain trials vs. baseline period exhibited exclusively lower connectivity scores for the pain periods, indicating a cortical decoupling during pain.

#### Negative Effects - connectivity

Throughout all pairs of cortical connections, we found a more positive relationship for pain intensity and connectivity than for pain unpleasantness and cortical connectivity (p<0.05, PALM corrected). As cortical processes disconnect during pain processing, the contrast between intensity and connectivity implies a stronger disconnection effect for unpleasantness (Figure 4, Supplementary Spreadsheet 1).

**Figure 4.**
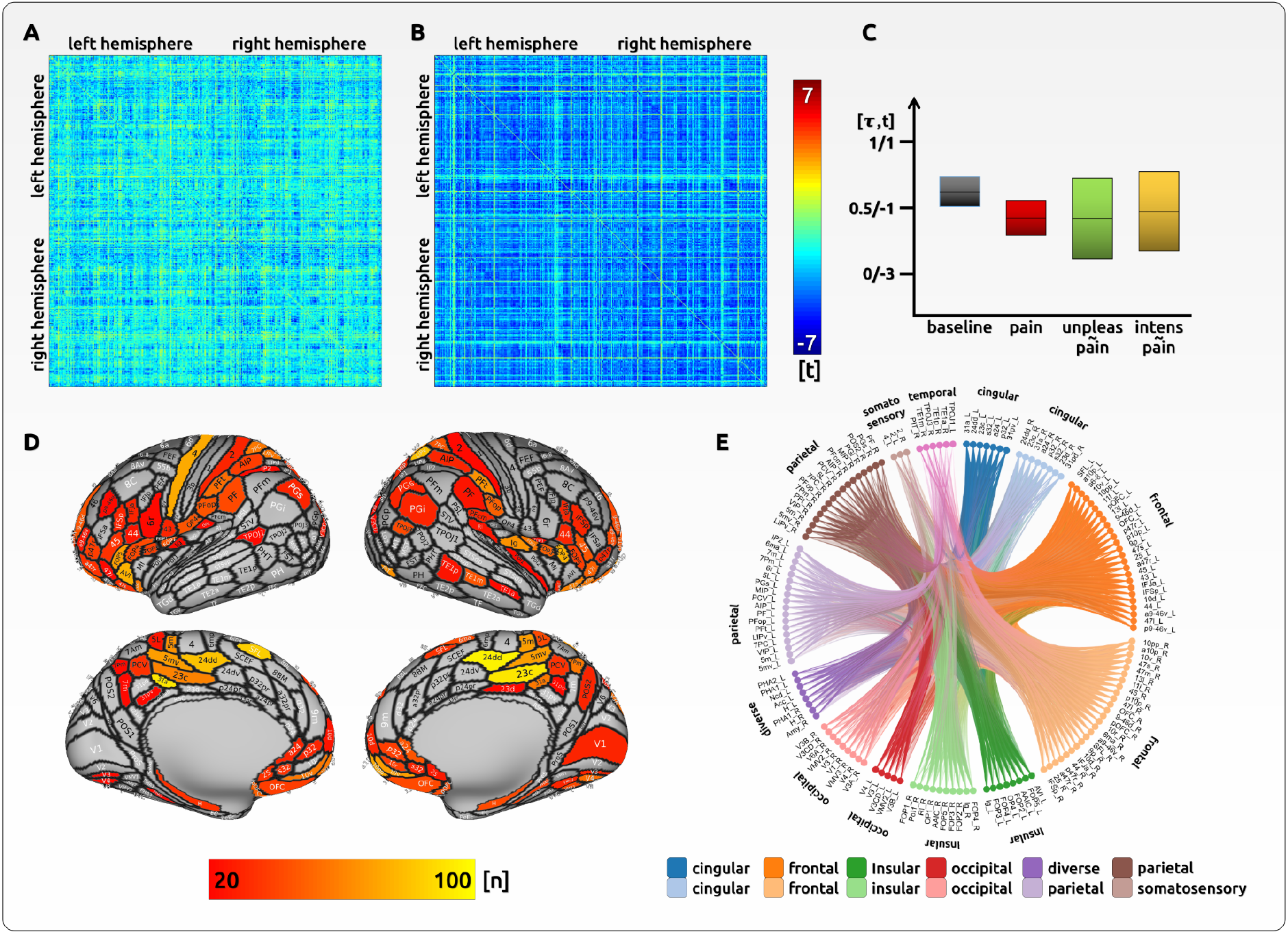
Differences between pain unpleasantness encoding and pain intensity encoding - connectivity. Several cortical connections reflect the differences between pain intensity and pain unpleasantness. (A) There is a general disruption of connectivity for pain trials compared to baseline periods (t-tests). (B) Single trial relationships with ratings show that the disruption of connectivity is better reflected by unpleasantness ratings (LME). (C) Averaged across all superthreshold regions: stronger connectivity for baseline periods (avg(τ)=0.43±0.13) than for pain trials (avg(τ)=0.36±0.11). T-values for pain encoding were lower for unpleasantness (avg(t)=-1.2±1.19) than for intensity (avg(t)=-1-18±1.19). (D) The figure shows the 3D map of regions that most frequently exhibit a significant effect for unpleasantness compared to intensity (threshold for presentation: n=20). (E) The circular plot shows regions that most frequently exhibit a significant effect for unpleasantness compared to intensity.

Taken together, the findings across BOLD and connectivity analyses exclusively indicate a stronger effect for unpleasantness. Areas that are *activated* through pain stimulation show a stronger *positive* relationship for unpleasantness. In a similar vein, stronger *negative* relationships in *deactivated* areas. In addition, unpleasantness ratings are related to a stronger cortical decoupling during pain processing. Consequently, the interpretation of the effects for activity and connectivity are identical: for all brain regions and cortical connections, cortical processes are always tighter connected to unpleasantness ratings. Please note that the differences are in a gradual fashion; due to the fact that both aspects of pain evaluation are highly correlated, we can not assume distinct processes for intensity and unpleasantness.

## 4. DISCUSSION

In the present study we investigated the cortical underpinnings of pain intensity and pain unpleasantness. Despite each aspect sharing the same fundamental core, we took advantage of the sufficient variability, which allowed us to investigate cortical functions related to the subjective experience of pain in a within-subject design at a single-trial level. We show that the emotional-affective aspect of pain is more deeply rooted in the human brain: throughout the brain, we revealed a stronger relationship for the pain-related cortical processes regarding the affective aspect compared to the sensory-discriminative aspect. No brain region showed a stronger relationship to pain intensity. As the processing of long-lasting pain is predominantly represented through disrupted connectivity, as shown here and in our previous work [14, 24], the current findings also point to a stronger disruption-related effect for pain unpleasantness than for pain intensity.

Regions that increased activity in response to pain had a more positive effect for unpleasantness, and regions and connections that exhibited decreased connectivity showed a more negative effect for unpleasantness. Consequently, unpleasantness ratings may reflect the more direct and intuitive processing of emotions in the human brain rather than the more complex and “minded” evaluation of pain intensity. Due to the methodological differences of previous work and their inherent weaknesses, the present study is not comparable to previous imaging studies [13, 25]. However, our results underline other research indicating the preferential processing of pain-related emotions in the human brain [26, 27]. It has been suggested that the feeling of unpleasantness belongs to the initial part of the noxious sensation and occurs before any conscious and cognitive processing of the sensation [28]. Therefore, the stronger effect of pain unpleasantness may better reflect the biological function of the pain system to prevent harm and to preserve the physical integrity of the body [29].

Indeed, there is a large framework of research that supports the notion that emotional processes are tightly associated with pain processing in the brain. Higher pain affect has been found to increase autonomic responses, such as skin conductance [30, 31], which in turn were found to encode pain-related cortical responses [32, 33]. Potentially emotion-related (“less minded”) autonomic responses were found to be more tightly related to cortical activity than ratings of pain intensity [32].

Our study complements previous findings by Zeidan and colleagues who shed light on individual abilities to modulate pain intensity and pain unpleasantness through meditation [25]. The better performing participants were able to make use of specific subunits of the pain modulation system. The participants who exhibited a better performance in the attenuation of pain unpleasantness, modulated the activity of the orbitofrontal cortex [25]. This region has been suggested to serve as a major hub of emotional modulation [34]. In a similar vein, the hub that modulates the ability to attenuate pain intensity through meditation could be located in the anterior insula [25]. Depending on the pain attenuation strategy, we would assume the existence of further cortical hubs that are differently utilised by individual subjects and which alter pain intensity and pain unpleasantness [14, 15].

### Conclusions

The present study shows that pain unpleasantness is more tightly related to cortical processing than the cognitive evaluation of pain intensity. These findings generalise to all brain regions that exhibit pain-related responses, irrespective of whether they increase or decrease their activity. However, the stronger relationship to the affective aspect does not imply that the assessment of pain intensity might be obsolete. Previous research has shown the usefulness of the assessment of intensity, which is more sensitive to change for certain pain conditions than unpleasantness [2, 8, 35]. Future studies need to shed further light on either aspect of pain evaluation under specific experimental (e.g. different types of applied pain) or disease-related conditions (episodic or chronic pain). However, across all conditions, we hypothesise temporal differences in the cascade of pain-related cortical processes: the cortical processes of pain unpleasantness evaluation are processed faster and may occur before the cognitive (“minded”) cortical processes of pain intensity evaluation.

## 5. DECLARATIONS

### Ethics approval and consent to participate

The study was approved by the Medical Sciences Interdivisional Research Ethics Committee of the University of Oxford (MSD-IDREC-C1-2014-157) and conducted in conformity with the Declaration of Helsinki.

### Consent for publication

Not applicable.

### Availability of data and materials

The data are available at the Open Science Framework (https://osf.io/tbc2u/). No custom code has been developed for this study.

### Authors’ contributions

A.S. and E.S. contributed to the conception and design of the study, acquisition and analysis of the data and drafting the manuscript and figures. A.M. contributed to the analysis of the data and drafting the manuscript. S.I. contributed to drafting the manuscript. V.W. contributed to the analysis of the data and drafting the manuscript.

## Acknowledgements

Not applicable.

