## Supplementary for "Pain and the Emotional Brain: Affective Rather than Cognitive Processes Drive the Cortical Encoding of Pain"

**Improved statistical modelling using a gradient vector**

For the BOLD data, we have pursued a further, more suitable approach, which we have moved to the supplement as it often did not pass reviewers' minds. Across all trials (modulated and unmodulated), we created a vector representing the single-trial difference between pain intensity and pain unpleasantness (UI_gradient). This gradient (example in Supplementary Table 2, in green) is better suited to distinguish the encoding of pain intensity and pain unpleasantness as it up-weights the larger differences between both pain descriptors. No differences are represented by zero. The conventional contrast ([-1 1], example in Supplementary Table 2, in orange) as being used in the main article would not differentiate between trials with large and no differences in the rating of both pain descriptors.
The gradient vector vector has been created for each trial as follows:

*(1) UI_gradient = [(U-I)/2; (I-U)/2]*

Furthermore, we have addressed potentially unexplained variance that may have an impact on the difference between intensity encoding and unpleasantness encoding. The variable "mean_rating" will control for a potential effect of the magnitude of pain ratings. The difference between intensity encoding and unpleasantness encoding might be influenced by the level of experienced pain (M=(U+I)/2). The variable "condition_mean" adds information on the average rating for each condition (reappraisal, counting, imagination, pain only), descriptor, and subject. The formula would be adapted as follows:

*(2) rating ~ fmri +* mean_rating + condition_mean + *UI_gradient:fmri + (1|subject)*

Naturally, as intensity and unpleasantness are highly correlated the variable "mean_rating" has a highly significant effect (t=16.95, p<0.001 for an right insular voxel). The "condition_mean" has no effect (all p>0.05). The results exhibit a better model fit compared to the model in the main article as represented by higher t-values. The statistical threshold is adjusted to account for multiple testings (p<0.05; PALM).

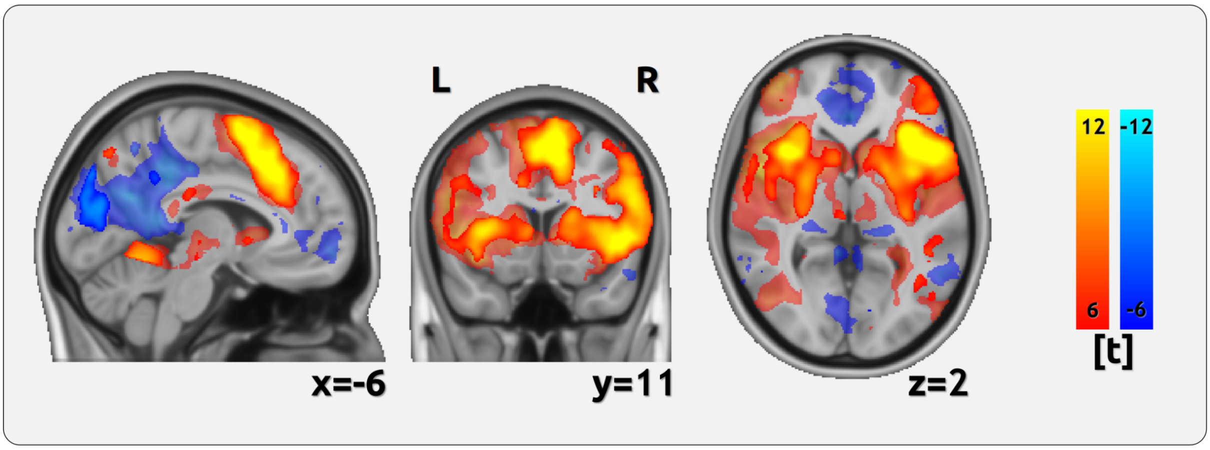


**Supplementary Figure 1 | Differences between pain unpleasantness encoding and pain intensity encoding.** Several cortical regions reflect the gradual differences between pain intensity and pain unpleasantness. Warm colours indicate regions that show a more positive relationship for pain unpleasantness; cold colours indicate brain regions that show a stronger negative relationship for pain unpleasantness. In order to better separate the different regions, we increased the statistical threshold beyond the significance level (t>4.9) for the positive effects. The pale colours indicate the contrast between pain trials and baseline (t>2).

**Results**

| max t-value | x | y | z | region |
|---|---|---|---|---|
| 13.5 | -45 | 14 | -2 | Inferior Frontal Gyrus |
| 13.5 | -9 | 14 | 47 | Paracingulate Gyrus |
| -9.13 | 2 | -89 | 24 | Precuneous Cortex |
| 13.4 | 35 | 15 | 5 | Insular Cortex |
| 11.6 | 26 | -55 | 47 | Superior Parietal Lobule |
| 11.4 | -43 | -40 | 48 | Superior Parietal Lobule |
| 10.1 | 37 | -24 | 54 | Precentral Gyrus |
| 10.3 | 48 | 5 | 35 | Precentral Gyrus |
| 11 | -10 | -53 | -12 | Lingual Gyrus |
| 9.5 | 43 | 29 | 31 | Middle Frontal Gyrus |
| -9.02 | 26 | 33 | 44 | Superior Frontal Gyrus |
| 10.2 | 29 | 51 | 15 | Frontal Pole |
| 9.73 | -33 | -18 | -2 | Insular Cortex |
| 9.98 | -60 | -19 | 26 | Postcentral Gyrus |
| 9.23 | -10 | -72 | 46 | Precuneous Cortex |
| -5.77 | 0 | 60 | -8 | Frontal Pole |
| -6.34 | 59 | -55 | 5 | Middle Temporal Gyrus |
| -5.96 | 38 | -72 | 27 | Lateral Occipital Cortex |
| -7.26 | 26 | -4 | -18 | Amygdala |

**Supplementary Table 1.** Differences in the encoding of pain intensity and pain unpleasantness. Positive effects indicate a stronger positive relationship for unpleasantness; negative relationships indicate a stronger negative relationship for unpleasantness (all p<0.05, PALM corrected).

**Data structure of one exemplary voxel.**

| subject | trial | rating | fMRI | rating_type | rating_code | gradient |
|---|---|---|---|---|---|---|
| 1 | 1 | 25 | 30.91 | intens | -1 | -2.5 |
| 1 | 1 | 30 | 30.91 | unpleas | 1 | 2.5 |
| 2 | 1 | 25 | 124.92 | intens | -1 | 5 |
| 2 | 1 | 15 | 124.92 | unpleas | 1 | -5 |
| 3 | 1 | 25 | 61.30 | intens | -1 | 7.5 |
| 3 | 1 | 10 | 61.30 | unpleas | 1 | -7.5 |
| 4 | 1 | 15 | 23.53 | intens | -1 | 2.5 |
| 4 | 1 | 10 | 23.53 | unpleas | 1 | -2.5 |
| 5 | 1 | 35 | 75.44 | intens | -1 | 5 |
| 5 | 1 | 25 | 75.44 | unpleas | 1 | -5 |
| 6 | 1 | 10 | 50.15 | intens | -1 | 2.5 |
| 6 | 1 | 5 | 50.15 | unpleas | 1 | -2.5 |
| 7 | 1 | 10 | 56.58 | intens | -1 | 2.5 |
| 7 | 1 | 5 | 56.58 | unpleas | 1 | -2.5 |
| 8 | 1 | 30 | 22.63 | intens | -1 | -7.5 |
| 8 | 1 | 45 | 22.63 | unpleas | 1 | 7.5 |
| 9 | 1 | 50 | 53.17 | intens | -1 | 0 |
| 9 | 1 | 50 | 53.17 | unpleas | 1 | 0 |
| ... | | | | | | |
| 11 | 44 | 35 | 57.02 | intens | -1 | 2.5 |
| 11 | 44 | 30 | 57.02 | unpleas | 1 | -2.5 |
| 12 | 44 | 10 | -53.56 | intens | -1 | 0 |
| 12 | 44 | 10 | -53.56 | unpleas | 1 | 0 |
| 13 | 44 | 35 | 120.45 | intens | -1 | 2.5 |
| 13 | 44 | 30 | 120.45 | unpleas | 1 | -2.5 |
| 14 | 44 | 45 | 40.20 | intens | -1 | 0 |
| 14 | 44 | 45 | 40.20 | unpleas | 1 | 0 |
| 15 | 44 | 40 | 143.39 | intens | -1 | 0 |
| 15 | 44 | 40 | 143.39 | unpleas | 1 | 0 |
| 16 | 44 | 30 | 41.96 | intens | -1 | 2.5 |
| 16 | 44 | 25 | 41.96 | unpleas | 1 | -2.5 |
| 17 | 44 | 50 | 85.72 | intens | -1 | -5 |
| 17 | 44 | 60 | 85.72 | unpleas | 1 | 5 |
| 18 | 44 | 55 | 181.06 | intens | -1 | -5 |
| 18 | 44 | 65 | 181.06 | unpleas | 1 | 5 |
| 19 | 44 | 35 | -10.43 | intens | -1 | -2.5 |
| 19 | 44 | 40 | -10.43 | unpleas | 1 | 2.5 |
| 20 | 44 | 20 | 3.32 | intens | -1 | 0 |
| 20 | 44 | 20 | 3.32 | unpleas | 1 | 0 |

**Supplementary Table 2 |** Please note the “mirrored” values of the gradient. It shows that the gradient approach is more suitable for the model as higher gradient values correspond with larger differences between the ratings of intensity and unpleasantness . This gradient score “weighs” the differences between both pain measures (compare trial 1 of subjects 1 and 2). The dichotomous model does not make any difference for the contrast of both measures. The dichotomous model compares all trials equally, even if there is no difference in rating for intensity and unpleasantness.
