## Supplementary for "Pain and the Emotional Brain: Affective Rather than Cognitive Processes Drive the Cortical Encoding of Pain"

(1) *Is it possible to set up a model to differentiate between the encoding of pain intensity and the encoding of pain unpleasantness?*

Based on previous submissions, this supplementary document can answer some questions that are usually not reported in the main manuscript text and can only implicitly be drawn from the main manuscript. We aim to give insight into the modelling for readers with less knowledge about linear mixed effects modelling. A wrong interpretation of the method can lead to incorrect interpretations of the findings.

(2) *Single trial ratings of intensity ratings have been acquired before ratings of unpleasantness.*

There might be an effect of order but we kept this sequence to ensure subjects did not confuse the rating process. Both descriptors rely on the same cortical data. In our view, an effect of rating order throughout all 44 trials appears unlikely. The magnitude of the ratings did not differ between intensity and unpleasantness.

(3) *Can we isolate one pain descriptor from the other, or can we only extract their difference?*

Both descriptors of pain are highly correlated with each other: there is no experience of high pain intensity without an experience of high pain unpleasantness. Previous experiments have aimed to independently modulate both aspects in different groups using hypnosis. The advantage of the present investigation is that we can distinguish between intensity and unpleasantness by relying on the very same single-trial cortical data. Hence, we can not claim that one or the other pain descriptor is associated with more or less cortical activity, because all ratings of intensity and unpleasantness are relying on the very same trial activation. Instead, we can draw conclusions on the question, which pain descriptor is more closely “correlated” to the underlying cortical data.
We have done that in the main text by a simple contrast [-1 1] and by a more complex model in the supplement by using a gradient vector. Both analyses point to a stronger effect for unpleasantness.

(4) *Do the obtained results support the conclusions that “the present study shows that pain unpleasantness is more tightly related to cortical processing than the cognitive evaluation of pain intensity.” Which brain regions reflect more the affective value of the pain, and which reflect the discriminative / sensory aspects of the stimuli?*

Yes, this can be easily drawn from the statistical model: The basis is a simple LME for the relationship between imaging data and pain perception at single trial level:

intensity ~ fmri + (1|subject)
unpleasantness ~ fmri + (1|subject)

The model can be independently applied to intensity as well as to unpleasantness. For the present investigation, in order to test whether one or the other aspect is tighter connected, we added a further factor: the “rating_type”. Through this contrast we can infer differences in the relationship between cortical data and pain perception. Please note the data structure in Supplementary Material.

Dichotomous model:
pain ~ fmri + rating_type:fmri + (1|subject)
Or
Gradient model:
pain ~ fmri + UI_gradient:fmri + (1|subject)

In simpler terms, do we find a stronger correlation with cortical data for intensity or for unpleasantness? The results of both models (dichotomous, gradient) show that the variance in the single trial cortical data are better explained by the variance of unpleasantness than the variance of intensity. All significant effects point to a tighter relationship between the affective ratings and brain activity. This applies to brain regions that exhibit *increased* activity to pain: there is a higher positive “correlation” for pain unpleasantness than for pain intensity. This also applies for the regions that exhibit *decreased* activity in response to pain: there is a stronger negative “correlation” for pain unpleasantness than for pain intensity. Any model that does not include a separating variable (gradient vector or dichotomous nominal variable) for the two descriptors of pain can not answer the difference in pain encoding.

(5) *Do the results only suggest a relationship between cortical responses and the difference between unpleasantness and intensity ratings?*

No. We tested the difference between the relationships. This can be drawn from the model formula.
Please also note that both rating scales are not directly comparable to each other. It is not clear whether the meaning of the lowest rating of intensity is equivalent to the lowest rating of unpleasantness. However, this is not needed for the model, where we test the differences in the relationship (the covariation) to the underlying cortical data.

(6) *Is it meaningful to present separate figures for the distinct analyses for intensity and unpleasantness?*

No. Both figures would largely show the same result as both descriptors of pain are highly correlated. In addition, as we were pooling the data across 4 conditions, the results would likely reflect the different pain states between conditions. In order to interpret the data, we have overlaid the encoding differences (the core of our analysis) on the contrast between all pain conditions with the pain-free periods during the experiment.

(7) *Are all data presented?*

Yes. For each voxel we found a stronger relationship between cortical data and pain unpleasantness compared to the relationship between cortical data and pain intensity. No voxel showed the opposite.

(8) *Do the statistical results show any differences between unpleasantness-related and intensity-related cerebral activation?*

No. Both descriptors of pain are relying on the same cortical data.

(9) *Do the t-maps represent cold pain-related cerebral activation?*

No. The t-maps of the current analysis represent the significant difference between fmri~unpleasantness and fmri~intensity. The pale maps in the figure represent the cold pain activation and deactivation. The pale maps have been included for a better interpretation of the findings.

(10) *Why do the activation clusters show a typical pattern of thermal pain-related cerebral activation as well as negative activation.*

These regions are known to be relevant for the processing of pain and should be activated/deactivated. The question is, whether the activation/deactivation in these regions is closer bound to unpleasantness or to intensity. Our model has answered this question in favour of unpleasantness.

(11) *How far do the findings go beyond what your group has already published?*

We used an existing data set for a new question, therefore the previous findings are unrelated to the new findings. The pain attenuation modulations in the previous data (Schulz et al., 2019) are statistically independent and irrelevant for the new research question. A further study (Schulz et al., 2020) uses functional connectivity metrics on pain attenuation, which again are not relevant here.

(12) *Shouldn't there be a clear and independent experimental dissociation between pain intensity and unpleasantness and an orthogonal manipulation of the two pain descriptors?*

If that would be correct, statistical inferences would be allowed to be drawn only from conditional differences in magnitudes. Here, we are testing whether both pain descriptors show a stronger relationship to the underlying cortical processes. The beauty and clear advantage of our approach is that both ratings rely on the very same cortical data. The linear mixed-effects model was provided with sufficient variance for intensity, unpleasantness, and their difference. Many biological and psychological variables are indeed of a gradual fashion (IQ, amount of attention, pain, hormone states). We often average and dichotomise these variables for simplicity but in many cases we would unnecessarily lose important variance in the data, as compared to single-trial analyses.

(13) *Does the experimental task imply that pain intensity ratings reflect purely cognitive processes while ratings of pain unpleasantness reflect purely affective processes.*

Of course not. Both ratings share the same fundamental core and are, hence, highly correlated. For this reason we do not make any statement e.g. on the cortical activity during the experience of pain unpleasantness vs. baseline.

*(14) Why can't we use a simpler contrast?*

Simpler contrasts like "rating_intensity ~ fmri", "rating_unpleasantness ~ fmri", or "all_ratings~fmri" would have estimated the relationship between cortical activity and pain ratings separately for each pain descriptor. However, this intermediate step is not meaningful for two reasons:
*First*, as intensity and unpleasantness are highly correlated, the map of 3D results would be very similar for both descriptors of pain.
*Second*, a separate analysis would be confounded by the 4 experimental conditions (unmodulated pain plus 3 pain attenuation interventions) because the unmodulated pain condition had higher pain ratings than the 3 pain-modulating conditions (unmodulated>reappraisal>safe place>counting). The results would reflect the differences across the conditions rather than the encoding of pain ratings. Furthermore, in order to prevent over-parameterisation, we did not include a random effect for slope.

This does not apply to the other included fixed effect (*rating_type:fmri*), which can be considered as a direct contrast between "pain_intensity~fmri" vs. "pain_unpleasantness~fmri". Both descriptors of pain rely on the same cortical data; the contrast reflects the encoding differences between intensity~fmri and unpleasantness~fmri irrespective of the experimental condition (which is identical). This aspect of the model formula essentially describes whether there is a stronger relationship for one descriptor compared to the other descriptor of pain.

*(15) Does the complex model contrast the predictive value of I and U on brain activity? And does the model show that some variance in brain activity evoked by pain stimuli is explained by the difference between U and I ratings.*

In simpler terms, we test whether there is a higher correlation between pain unpleasantness and brain activity than between pain intensity and brain activity.
That also means that the trial-by-trial variation of brain activity is best explained by the unpleasantness vector. The gradient vector (the difference between I and U) weights the encoding difference as an advantage compared to a more simple categorical contrast (see Supplementary Material). That means that the unpleasantness ratings are better suitable to predict brain activity (and vice versa).

*(16) Does the model show that some variance in brain activity evoked by pain stimuli is explained by the difference between U and I ratings.*

No. This suggested question would require the following model. In our view, this model would not make much sense.

fMRI~UI_gradient + (1|subject)

(17) *Can we separately investigate both descriptors of pain experience through an independent experimental variation of intensity and unpleasantness*.

This would be very desirable, indeed. In our view, however, under normal circumstances there will always be a tight correlation between ratings of pain intensity and ratings of pain unpleasantness. A previous study series attempted a dissociation by hypnotising participants to independently variate pain intensity and pain unpleasantness. The researchers kept one descriptor (e.g. intensity) constant and modulated the other descriptor (e.g. unpleasantness).

However, this approach has two important limitations. First, unlike our study, the cortical data were acquired in different sessions with probably different participants. This introduces between-subject or between session variance into the statistical analysis. In our study pain intensity and pain unpleasantness were relying on the very same cortical data. This issue is noticeable for the activity in the posterior part of the anterior cingulate cortex which is slightly shifted in both publications [1, 2]. Further issues with the attempt to independently modulate both descriptors of pain have been discussed in the main article.

In addition, an independent separation of both pain descriptors is not needed as we can rely on sufficient variation between intensity and unpleasantness (75% distinct ratings).

*(18) Why does the gradient regressor consist of perfectly anticorrelated parts: [U-I/2] and [i-U/2]?*

Indeed, the correlation coefficient between *[U-I/2] and [I-U/2]* is exactly - 1. This may sound complicated and is the reason we moved the actually more appropriate analysis with the gradients to the supplement. A look at Table 2 in the supplement may help here. Each trial has a double entry of fMRI values as each trial has two ratings: intensity and unpleasantness. In simple terms, the contrast model uses the coding [-1 1] to compare unpleas~fmri vs. intensity~fmri. In the more complex model, we have weighted each trial according to the distance of both pain descriptors, e.g. [-2.5 2.5]. Similar to the contrast coding [-1 1], the gradient coding [-2.5 2.5] has mirrored entries, too.
